# Molecular basis of autoimmune disease protection by MDA5 variants

**DOI:** 10.1101/2024.10.03.616466

**Authors:** Rahul Singh, Alba Herrero del Valle, Joe D. Joiner, Marleen Zwaagstra, Brian J. Ferguson, Frank J. M. van Kuppeveld, Yorgo Modis

## Abstract

MDA5 recognizes dsRNA from viruses and retroelements. Cooperative filament formation and ATP-dependent proofreading confer MDA5 with the necessary sensitivity and specificity for dsRNA. The gene encoding MDA5 is a hotspot for disease-associated variants. Many MDA5 variants are associated with protection from autoimmune disease, while increasing the risk of infection and chronic inflammation, but how these variants affect MDA5-dependent RNA sensing remains unclear. Here, we determine the consequences of autoimmune-protective MDA5 variants on the molecular structure and activities of MDA5. The rare variants E627* and I923V reduce the cellular interferon response to picornavirus infection and are deficient in filament formation. The I923V variant is ATPase hyperactive, causing premature dissociation from dsRNA. Cryo-EM structures of MDA5 I923V bound to dsRNA at different stages of ATP hydrolysis reveal smaller RNA binding interfaces, leading to excessive proofreading activity. Variants R843H and T946A, which are genetically linked and cause mild phenotypes, have no effect on dsRNA recognition, suggesting an indirect disease mechanism. We conclude that the autoimmune-protective MDA5 variants lead to a loss of MDA5-dependent signaling via multiple distinct mechanisms.

## Introduction

Viruses deliver or generate RNA in the cytosol. Cytosolic dsRNA, one of the most proinflammatory molecular signals from viruses and retroelements (Hur, 2019), is sensed in vertebrates by RIG-I (Kato *et al*, 2005; Yoneyama *et al*, 2005), MDA5 (Kato *et al*, 2008; Kato *et al*, 2006; Yoneyama *et al*., 2005), LGP2 (Li *et al*, 2009; Pippig *et al*, 2009; Satoh *et al*, 2010; Uchikawa *et al*, 2016; Venkataraman *et al*, 2007), and protein kinase R (Kumar *et al*, 1994; Vijay-Kumar *et al*, 2005). These are complemented in mammals by the oligoadenylate synthases (Chebath *et al*, 1987; Kristiansen *et al*, 2010), ZBP1, and in humans, NLRP1 (Bauernfried *et al*, 2021). MDA5 recognizes dsRNAs longer than 100 bp (Kato *et al*., 2008; Kato *et al*., 2006) and is the primary innate immune sensor for many viruses, including SARS-CoV-2 (Rebendenne *et al*, 2021; Yin *et al*, 2021). MDA5 is a superfamily 2 helicase with two RecA-like domains (Hel1 and Hel2), an insert domain (Hel2i), and a C-terminal domain (CTD) linked to Hel2 by a pair of α-helices known as the pincer domain (**Figure 1A**). These domains cooperatively bind dsRNA to form helical MDA5-dsRNA filaments (Berke & Modis, 2012; Berke *et al*, 2012; Peisley *et al*, 2011; Wu *et al*, 2013; Yu *et al*, 2021; Yu *et al*, 2018). Filament formation induces the tandem N-terminal caspase recruitment domains (CARDs) of MDA5 to oligomerize (Berke & Modis, 2012; Peisley *et al*., 2011). MDA5 CARD oligomers recruit MAVS via CARD-CARD interactions, nucleating assembly of MAVS CARD microfibrils (Hou *et al*, 2011; Wu *et al*, 2014), which function as supramolecular organizing centers for downstream effectors of the interferon-β (IFN-β) and NF-κB inflammatory responses (Hou *et al*., 2011; Kagan *et al*, 2014; Kato *et al*., 2008; Kato *et al*., 2006).

**Fig. 1.**
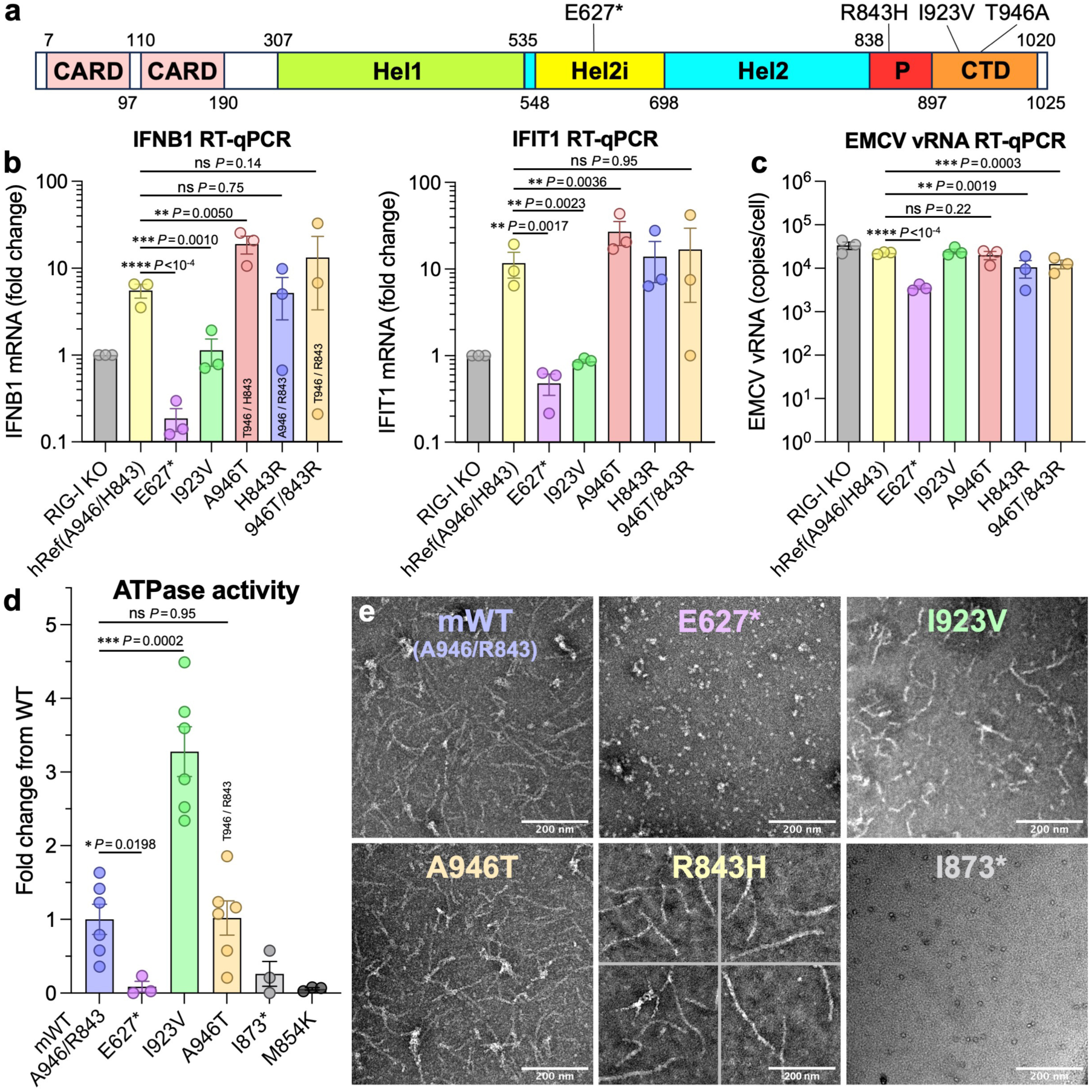
Effects of T1D-protective MDA5 variants on the antiviral interferon response and the ATPase and filament forming activities of MDA5. (**A**) MDA5 domain organization. CARD, caspase recruitment domain; Hel1 and Hel2, RecA-like helicase domains; Hel2i, Hel2 insert domain; P, pincer domain; CTD, C-terminal domain. (**B**) RT-qPCR quantification of *IFNB1* and *IFIT1* mRNA in A549 RIG-I KO cells stably expressing the indicated MDA5 variant under a doxycycline-inducible promoter 7 h after infection with encephalomyocarditis virus (EMCV). I923V and E627* inhibit the antiviral response. hRef, human reference sequence. (**C**) RT-qPCR quantification of EMCV RNA 7 h post-infection. (**D**) ATPase activities of T1D-protective MDA5 variants, normalized to WT and with zero-fold change set to the ATPase activity of the M854K variant (t = 3 min). Error bars represent mean ± SEM (3 or 6 measurements from 1 or 2 independent experiments, respectively). Source data for panels *(B-D)* are provided as a Source Data file (**Data S1**). mWT, mouse wild-type. (**E**) Negative-stain electron micrographs of MDA5 variants with 1-kb dsRNA and 4 mM ATP. Scale bars, 200 nm. Each micrograph is representative of at least six images.

Viral dsRNA can be difficult to distinguish from endogenous RNA. Innate immune responses must be sensitive enough to detect infection and specific enough to avoid activation by cellular RNA. The ATPase activity of MDA5 confers the necessary specificity of dsRNA recognition. Conformational changes coupled to ATP hydrolysis fulfill a proofreading function by promoting dissociation of MDA5 from endogenous dsRΝΑs (Yu *et al*., 2021; Yu *et al*., 2018), which are shorter and have weaker base-pairing due to mismatches and A-to-I deamination by ADAR1 (Ahmad *et al*, 2018; Chung *et al*, 2018; Liddicoat *et al*, 2015). The cooperativity of both filament formation and ATP hydrolysis by MDA5 confers sensitivity by encoding greater stability for long MDA5 filaments such that only filaments formed on longer dsRNAs of viral origin persist long enough to activate signaling (Singh *et al*, 2024; Yu *et al*., 2018).

The gene encoding MDA5, *IFIH1*, is a hotspot for natural variants with diverse clinical associations. Approximately 40 missense variants are associated with autoinflammatory disease, including Aicardi-Goutières syndrome (AGS) and Singleton-Merten syndrome (SMS) (Rice *et al*, 2014; Rice *et al*, 2020; Rodero & Crow, 2016; Rutsch *et al*, 2015). In most of these variants, the amino acid substitutions inhibit ATP hydrolysis, either directly (e.g. R337G) (Rice *et al*., 2014) or allosterically (e.g. M854K) (Yu *et al*., 2021). This disrupts ATP-dependent proofreading and allows MDA5 signaling complexes to form on endogenous dsRNAs (Garau *et al*, 2019; Rice *et al*., 2014; Rice *et al*., 2020; Takeichi *et al*, 2018; Yu *et al*., 2021). Other autoinflammatory variants map to the RNA binding interface and promote signaling from endogenous RNAs by increasing the RNA binding affinity of MDA5 (Rice *et al*., 2014). A distinct set of variants reduce the risk of developing certain autoimmune diseases, most notably type 1 diabetes (T1D). Missense single-nucleotide polymorphisms (SNPs) resulting in the MDA5 variants E627* (rs35744605), R843H (rs3747517), I923V (rs35667974), and T946A (rs1990760) are associated with protection from T1D (de Azevedo Silva *et al*, 2015; Gorman *et al*, 2017; Jermendy *et al*, 2018; Liu *et al*, 2009; Nejentsev *et al*, 2009; Smyth *et al*, 2006; Vasseur *et al*, 2011). The E627* and I923V variants are rare while R843H and T946A are common. The T946A variant results from an A:T to G:C base pair substitution. The T1D-protective G:C (Ala946) allele frequency is 30-50% in Caucasians, and 70-80% Africans and Asians (de Azevedo Silva *et al*., 2015; Gorman *et al*., 2017; Liu *et al*., 2009; Nejentsev *et al*., 2009; Vasseur *et al*., 2011). Similarly, R843H results from a G:C to A:T base pair substitution and the A:T (H843) allele frequency is 30-40% in Caucasians and Africans, and 70% in Asians (de Azevedo Silva *et al*., 2015; Gorman *et al*., 2017; Liu *et al*., 2009; Nejentsev *et al*., 2009; Vasseur *et al*., 2011). Variants T946A and R843H are in strong linkage disequilibrium with each other (Liu *et al*., 2009; Nejentsev *et al*., 2009; Pedergnana *et al*, 2016) such that Ala946 is predominantly found with His843, and Thr946 with Arg843 (Gorman *et al*., 2017; Pedergnana *et al*., 2016; Vergara *et al*, 2016). We note that most human *IFIH1* reference sequences contain the alleles encoding Ala946/His843. Wild-type (WT) mouse MDA5 contains Ala946/Arg843.

Whereas the autoinflammatory MDA5 variants increase basal IFN-β signaling, T1D protection correlates with reduced MDA5-dependent signaling (Gorman *et al*., 2017; Hoffmann *et al*, 2015; Shigemoto *et al*, 2009). MDA5 knockout (KO) nonobese diabetic (NOD) mice are fully protected from T1D-like disease, and heterozygous (MDA5^+/−^) mice expressing half of the wild-type level of MDA5 are significantly protected from disease (Lincez *et al*, 2015). The E627* and I923V variants decreased IFN-β signaling in a cell-based luciferase reporter assay (Shigemoto *et al*., 2009). Basal IFN-β transcription was slightly reduced in the T946A variant in human peripheral blood mononuclear cells (PBMCs) (Gorman *et al*., 2017), transfected human or mouse cells (Gorman *et al*., 2017; Hoffmann *et al*., 2015; Shigemoto *et al*., 2009), and mice(Gorman *et al*., 2017). The R843H variant had no effect on IFN-β transcription (Shigemoto *et al*., 2009). This suggests that variants E627*, I923V, and T946A either reduce the intracellular MDA5 protein concentration or alter the biochemical properties of the protein. There have been conflicting reports regarding whether these variants are transcribed at different levels but overall there is no compelling evidence of statistically significant differences in transcript or protein concentration for any MDA5 missense variant (Downes *et al*, 2010; Gorman *et al*., 2017; Liu *et al*., 2009; Marinou *et al*, 2007; Shigemoto *et al*., 2009; Zouk *et al*, 2014). Hence, how the T1D-protective MDA5 variants alter dsRNA sensing by MDA5 remains unknown. However, there is a strong clinical link between T1D onset and recent infection with RNA viruses, in particular coxsackieviruses and other enteroviruses. T1D patients have more frequent enterovirus infections, which precede the appearance of prediabetic markers, including autoantibodies (Hyoty & Taylor, 2002). MDA5 recognizes RNA from *Picornaviridae*, including enteroviruses (Kato *et al*., 2006; Wang *et al*, 2010), which have evolved mechanisms to suppress IFN-β transcription (Hato *et al*, 2007; Visser *et al*, 2019). MDA5-induced inflammation and cell death in the pancreas following rotavirus infection contributes to autoimmune destruction of pancreatic β-cells (Dou *et al*, 2017). Conversely, MDA5 KO mice are protected from disease upon infection with a β-cell-tropic coxsackievirus (Lincez *et al*, 2021). Therefore, a plausible hypothesis is that MDA5-dependent interferon production and inflammation following viral infection can trigger autoimmune β-cell killing.

Here, we examine the consequences of T1D-protective MDA5 substitutions on the structure and activities of MDA5. We show that variants E627*, I923V, and T946A reduce the IFN-β response to picornavirus infection. The E627* variant cannot bind RNA or form filaments. The I923V variant has increased ATPase activity, and reduced filament stability. Cryo-EM structures of the I923V variant bound to dsRNA at different stages of ATP hydrolysis reveal that the Ile923 sidechain regulates the conformational changes necessary for ATP hydrolysis. The T946A substitution does not affect RNA recognition, suggesting an indirect T1D protection mechanism. Hence, we have uncovered multiple, loss-of-function pathways that lead to T1D protection via distinct molecular mechanisms.

## Results

### T1D-protective MDA5 variants impair type I interferon response to picornavirus infection

Most but not all previous studies report that T1D-protective MDA5 variants reduce interferon responses. Transient overexpression of the E627* and I923V variants in human or mouse cells reduced *IFNB1* and *IFIT1* transcription with or without dsRNA (poly(I:C)) stimulation (Gorman *et al*., 2017; Hoffmann *et al*., 2015; Shigemoto *et al*., 2009), or upon picornavirus infection (Dou *et al*., 2017; Gorman *et al*., 2017), resulting in increased viral loads (Dou *et al*., 2017). Similarly, the T946A substitution reduced transcription of type I and type III interferon signature genes in transiently transfected HEK293T cells (Gorman *et al*., 2017; Hoffmann *et al*., 2015) and in knock-in mice (Gorman *et al*., 2017). The T946A substitution also promoted picornavirus replication and increased infection mortality in mice (Gorman *et al*., 2017). However, the T946A substitution was reported in other studies to increase *IFNB1* and *IFIT1* transcription and reduce picornavirus replication in transiently transfected HEK293 cells (Dou *et al*., 2017), and to increase IFN-λ transcription in human pancreatic islets infected with coxsackievirus (Domsgen *et al*, 2016). In a further study, the T946A substitution had no effect on *IFNB1* transcription in transiently transfected mouse embryonic fibroblasts (Shigemoto *et al*., 2009). Regarding the R843H variant, most studies conclude that its association with T1D protection is explained by its co-occurrence in most human subjects with the T946A variant (Gorman *et al*., 2017; Nejentsev *et al*., 2009; Shigemoto *et al*., 2009), but one study reported that the His843/Thr946 variant (which rarely occurs in humans) increased type I and type III interferon transcription in HEK293 cells (Hoffmann *et al*., 2015). To clarify the effects of these T1D-protective variants on MDA5-dependent interferon signaling, we generated RIG-I KO A549 human lung epithelial cell lines stably expressing each MDA5 variant under a doxycycline-inducible promoter. RIG-I KO A549 cells have very little, if any, endogenous MDA5 activity based on the absence of poly(I:C)-induced *IFNB1* transcription in these cells (Teague *et al*, 2024). Cells were infected with encephalomyocarditis virus (EMCV), a model picornavirus. We used a recombinant EMCV containing mutations in its Leader protein, i.e. the viral IFN antagonist protein (Hato *et al*., 2007), to achieve strong type I IFN transcription signals. *IFNB1* and *IFIT1* transcription as well as EMCV RNA replication were quantified by RT-qPCR seven hours after infection. We found that the E627* and I923V variants both failed to induce a type I interferon response (**Figure 1B** and **Figure S1**). Cells expressing I923V MDA5 had the same *IFNB1* and *IFIT1* transcription levels as the untransduced reference cell line. The E627* variant showed a further reduction in *IFNB1* transcription, and to a lesser extent *IFIT1* transcription, below the levels in the reference cell line. This was not due to increased cell death as transcription levels were normalized to *Actin* expression. The reference His843/Ala946 (R843H/T946A) variant induced *IFNB1* and *IFIT1* transcription to the same extent as the common Arg843/Thr946 (A946T) variant and the rare Arg843/Ala946 (H843R) variant, but the rare His843/Thr946 variant increased *IFNB1* and *IFIT1* transcription (**Figure 1B**), consistent with a previous study (Hoffmann *et al*., 2015). None of the variants significantly altered baseline *IFNB1* or *IFIT1* transcription in the absence of EMCV infection (**Figure S1**). EMCV replicated efficiently in all cell lines (**Figure 1C**). Hence, the effects of the variants on *IFNB1* and *IFIT1* transcription were not due to differences in viral RNA replication or viral dsRNA availability. In summary, our data show that the rare T1D-protective E627* and I923V variants cause loss of MDA5-dependent IFN-β signaling, in agreement with previous studies, whereas the protective R843H and T946A variants had no significant effect on EMCV-induced signaling in the combination commonly found in human subjects (R843H/T946A).

### ATPase and filament forming activities of T1D-protective MDA5 variants

To assess the effects of T1D-protective substitutions on the biochemical properties of MDA5, we purified recombinant mouse WT, E627*, I923V, and A946T MDA5 proteins. The E627* protein lacked ATPase activity (**Figure 1D**) and did not form filaments on 1 kb dsRNA, based on negative stain electron microscopy imaging (**Figure 1E**). In stark contrast, the I923V variant had 2.7-fold higher ATPase activity than WT MDA5 (**Figure 1D**) and formed filaments on dsRNA, although the filaments were shorter than those formed by WT MDA5 (**Figure 1E**). The A946T variant was notable in that its ATPase and filament forming activities were the same as WT MDA5 (**Figure 1D,E**). We note that the A946T variant was previously reported to have reduced ATPase activity (Funabiki *et al*, 2014) but we observed no statistically significant difference in the ATPase activities of A946T and WT purified recombinant mouse MDA5. We conclude that T1D-protective substitutions have pleiotropic effects on the biochemical activities of MDA5 and therefore act via distinct mechanisms. E627* is a simple loss of function mutant that lacks signaling activity because it cannot bind dsRNA. We note that truncation of the CTD (I873*) was sufficient to abrogate both ATPase activity and filament formation on dsRNA (**Figure 1D,E**). The loss of signaling activity of the I923V variant is explained by its increased ATPase activity, because ATP hydrolysis promotes dissociation from dsRNA (Yu *et al*., 2021; Yu *et al*., 2018). The increased ATPase activity of the I923V variant will therefore reduce the stability of signaling complexes, as evidenced by the reduced I923V MDA5-dsRNA filament length. This is the converse of substitutions that inhibit ATP hydrolysis, which reduce dissociation of MDA5 signaling complexes from dsRNA, including endogenous dsRNAs, and hence trigger autoinflammatory signaling (Rice *et al*., 2014; Yu *et al*., 2021).

### MDA5 variant I923V has reduced affinity for dsRNA

The reduced number and average length of filaments formed by the I923V variant on dsRNA (**Figure 1E**) suggests that the I923V substitution reduces the net dsRNA binding affinity of MDA5. To quantify this effect, we used differential scanning fluorimetry (DSF) and bio-layer interferometry (BLI) with purified recombinant mouse MDA5 proteins. The WT, I923V, A946T, and R843H variants had the same thermostability based on their DSF melting curves. These variants were significantly more thermostable under filament forming conditions than as monomers in solution, with the melting temperature increasing by 9-10°C following addition of 1-kb dsRNA (**Figure 2A,B**). In a more direct measure of binding affinity, BLI measurements showed that the I923V variant had a significantly lower binding affinity for 200-bp and 300-bp dsRNA than WT MDA5. The overall dissociation constants (*K*_d_ values) of MDA5 I923V for 200-bp RNA were 5.6 nM and 1.3 nM, respectively (**Figure 2C,D** and **Figure S2**). In contrast, the BLI binding curves for the A946T and R843H variants were the same as WT MDA5, within experimental error, with 200-bp, 300-bp dsRNA (**Figure 2C**). In BLI measurements with 1-kb dsRNA, the I923V, A964T and R843H variants all bound similarly to WT MDA5, but *K*_d_ values could not be accurately determined due to the complex shape of the BLI curves early in the binding step (**Figure 2C** and **Figure S2**). We conclude that the I923V substitution reduces the affinity of MDA5 for dsRNA, whereas the A946T and R843H substitutions have no significant effect on dsRNA binding affinity and kinetics.

**Fig. 2.**
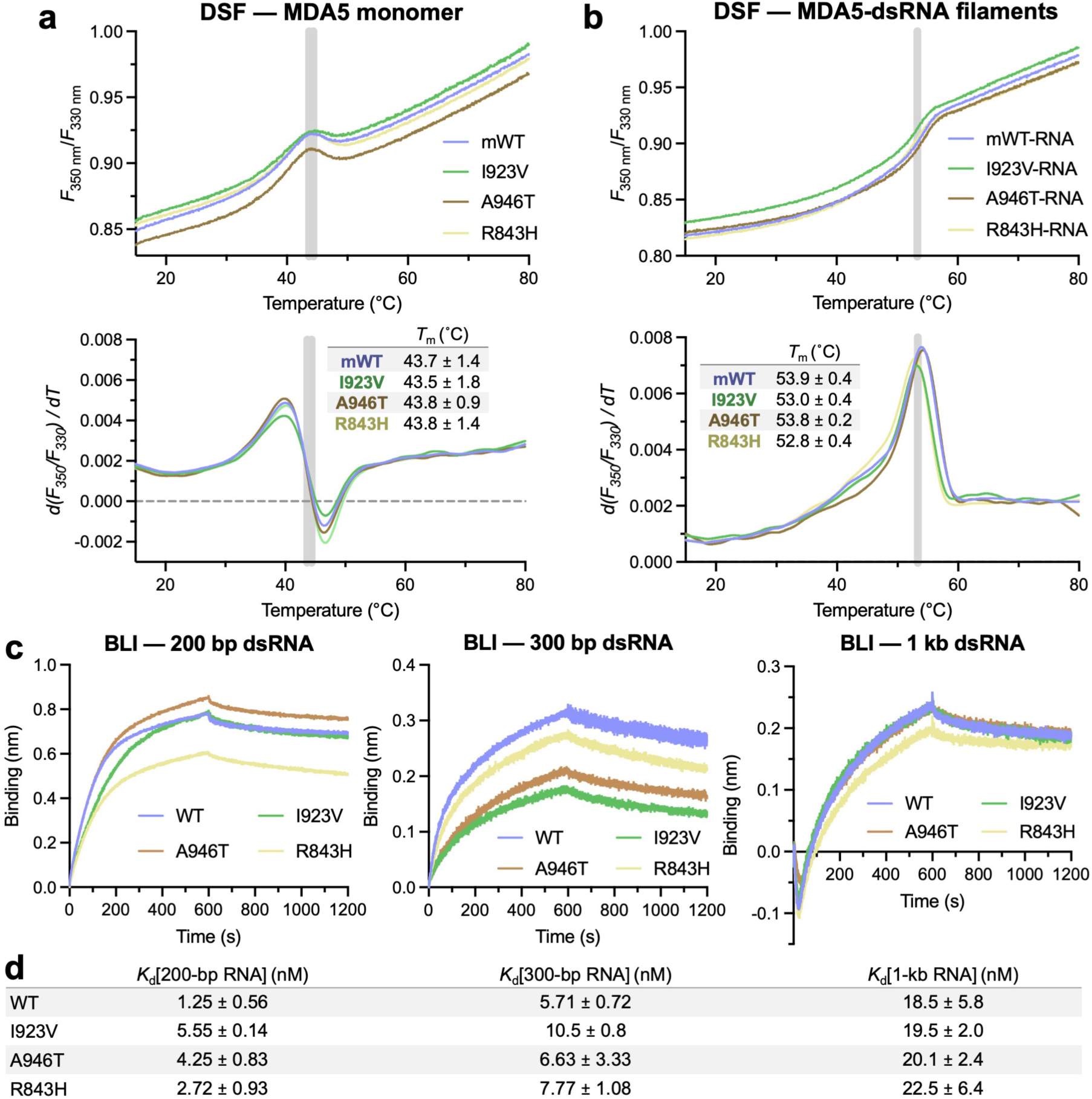
Thermostability and RNA binding affinities of T1D-protective MDA5 variants. (**A**) Differential scanning fluorimetry (DSF) of WT and variant MDA5 proteins. Intrinsic protein fluorescence at 330 nm and 350 nm was measured and the fluorescence ratio plotted as a function of temperature. Grey lines indicate the melting temperatures (*T*_m_) of the variants. (**B**) The *T*_m_ of the MDA5 variants was higher in the presence of 1-kb dsRNA. (**C**) Bio-layer interferometry (BLI) with 3’-biotinylated dsRNA immobilized on a streptavidin sensor and 125 nM mouse MDA5 in the mobile phase (see also **Figure S2**). The I923V variant has a slightly lower affinity for 200-bp and 300-bp dsRNA. (**D**) Dissociation constants (*K*_d_ values) derived from the curves in (*C*). The uncertainties are the s.e.m. from three independent experiments. Curves from a single representative experiment are shown in (*A-C*). For 1-kb RNA, data from the first 100 s were omitted for *K*_d_ calculation because the curve shape in that time interval was complex and could not be fitted.

### I923V MDA5 has an altered distribution of conformational states during catalysis

Structural studies have shown that MDA5 variants associated with autoinflammatory disease impair discrimination of endogenous RNA from viral RNA by altering the conformational changes that are coupled to the ATPase cycle, directly or indirectly (Yu *et al*., 2021). To gain a mechanistic understanding of the effects of the individual I923V and A946T substitutions on MDA5 function, we determined cryo-EM structures of mouse MDA5-dsRNA filaments containing each of these substitutions at different stages of ATP hydrolysis (**Figure 3A**). Four cryo-EM datasets were collected: I923V MDA5 filaments with ATP, transition state analog ADP-AlF_4_, or no nucleotide bound, and A946T filaments without nucleotide (**Table S1** and **Figure S3**). 3-D classification of the maps with helical symmetry averaging applied allowed us to analyze the helical twist distributions of the filament segments for each dataset. We found that on binding ATP, filaments formed from the I923V variant mostly had an intermediate helical twist (81°-91°) instead of the low twist (71°-81°) observed in ATP-bound WT MDA5 filaments (**Figure 3B**). Moreover, with ADP-AlF_4_ bound, the I923V MDA5 filaments had a broad distribution of low to intermediate twists instead of the narrow distribution of intermediate twists observed in the transition state of WT MDA5. In the absence of nucleotide, the I923V and A946T variants both had similar twist distributions to WT MDA5, with intermediate to high twists (81°-96°; **Figure 3B**). Hence, the I923V substitution alters the helical twist distributions of MDA5-dsRNA filaments in the ATP-bound and ADP-AlF_4_-bound states.

**Fig. 3.**
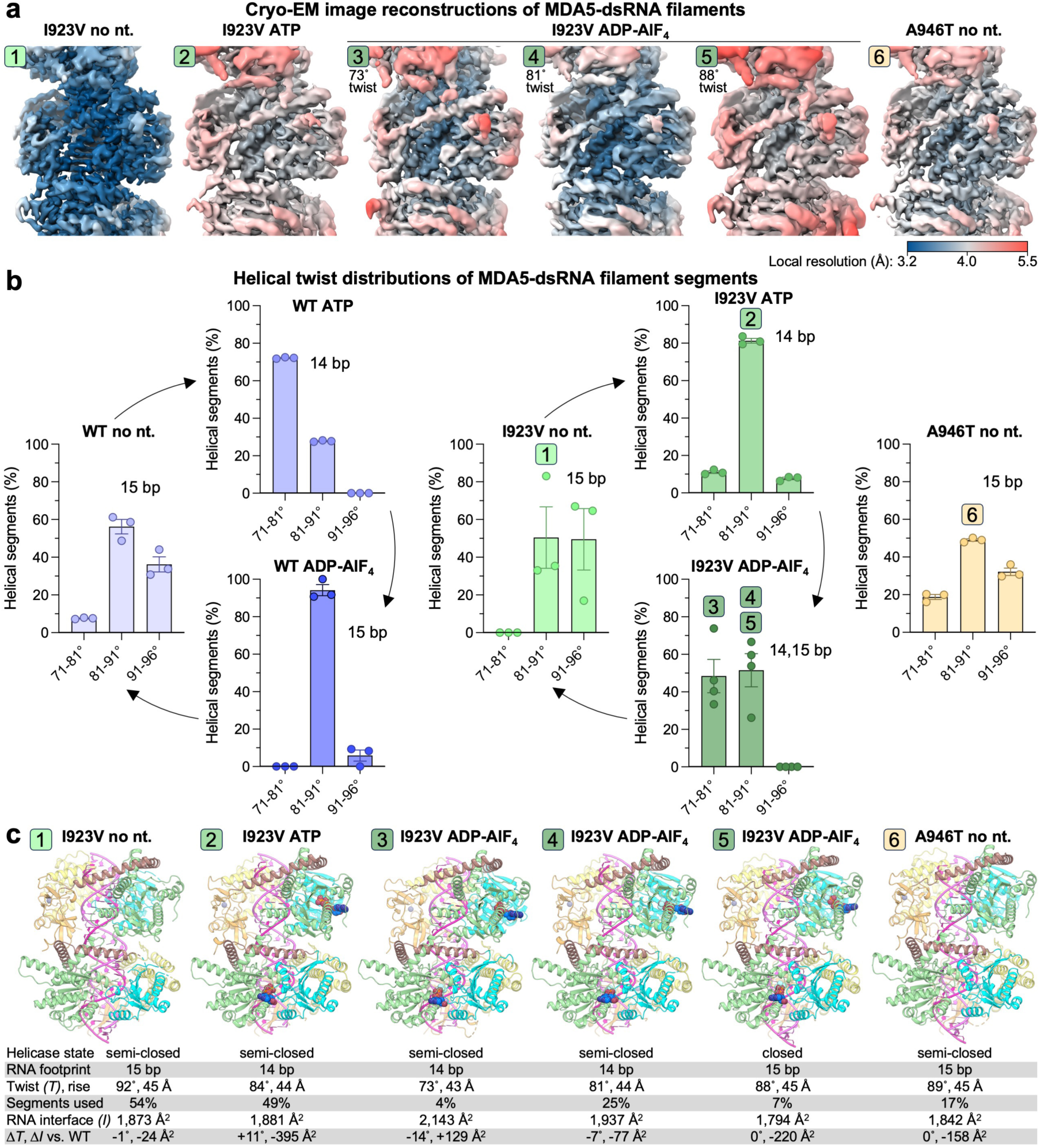
Cryo-EM structures of I923V and A946T MDA5 bound to dsRNA at different stages of ATP hydrolysis. (**A**) Cryo-EM image reconstructions of MDA5-dsRNA filaments with helical symmetry averaging. (**B**) Helical twist distributions of MDA5-dsRNA filament segments after 3-D classification. (**C**) Atomic models and their structural parameters. Two helical subunits are shown for each model. See **Figure S3** for Fourier Shell Correlation curves.

Processing and refinement of our four cryo-EM datasets yielded six maps with resolutions sufficient to build and refine atomic models of the protein, dsRNA, and bound nucleotides. The ATP-bound I923V dataset and nucleotide-free I923V and A946T datasets each yielded one map, whereas the ADP-AlF_4_-bound I923V dataset yielded three different maps (**Figure 3A**). The atomic models of the nucleotide-free states of the I923V and A946T variants are similar to previously reported nucleotide-free structures of WT MDA5 (Yu *et al*., 2018), with 15-bp RNA footprints, intermediate to high helical twists (89°-92°), and protein-RNA interaction areas of 1,800 to 1,900 Å^2^ per MDA5 subunit calculated with PISA (Krissinel & Henrick, 2007) (**Figure 3C**). The ATP-bound I923V structure has the same 14-bp RNA footprint as ATP-bound WT MDA5 (Yu *et al*., 2018), but a significantly higher helical twist (84°, versus 73° for WT) and a 20% smaller protein-RNA interaction area (**Figure 3C**). The three ADP-AlF_4_-bound I923V structures have helical twists of 73°, 81°, and 88°, respectively, reflecting the helical twist distribution of ADP-AlF_4_-bound I923V filaments (**Figure 3B**). The 73°-twist and 81°-twist structures have 14-bp RNA footprints whilst the 88°-twist structure has a 15-bp footprint. The 88°-twist structure is similar to the ADP-AlF_4_-bound WT MDA5 structure, albeit with a 10% smaller protein-RNA interaction area (**Figure 3C**). However, the 73°-twist and 81°-twist structures both differ from the WT transition state structure in that they retain the same 14-bp footprint and lower twist as the ATP-bound ground state. Additionally, only in the 88°-twist structure are the Hel1 and Hel2 domains in the catalytically competent closed conformation (as defined based on the Hel1-Hel2 rotation angle – see Materials and Methods). Indeed, the 73°-twist and 81°-twist structures are in the semi-closed conformation in which the Hel2 domain is not engaged with the nucleotide. Together, the structural features of the 73°-twist and 81°-twist ADP-AlF_4_-bound I923V structures indicate they do not represent the catalytic transition state but rather intermediates that more closely resemble the ground state. In summary, the I923V and A946T variants adopt similar sets of structures as WT MDA5 but the I923V variant has a smaller protein-RNA interaction area than WT in the ATP-bound state, and I923V can accommodate ADP-AlF_4_ in its active site in a ground state-like conformation with a 14-bp RNA footprint and as well as in the closed transition state with a 15-bp footprint.

### Isoleucine is required at position 923 to sterically regulate ATPase activity

Closer examination of the structures of the I923V and A946T variants did not reveal any noteworthy changes in the fold of the CTD, where both substitutions are located (**Figure 4A,B**). In all available structures, the side chain of residue 923 contributes to the hydrophobic core of the CTD and forms multiple van der Waals contacts with surrounding side chains. In the structures of the I923V variant, there are minor differences in the side chain positions of several nearby residues occur, including Glu924, Lys925, His974 and Tyr1015, possibly to compensate for the smaller size of the valine side chain in the hydrophobic core (**Figure 4C,D**). As a result, a hydrogen bond between the side chain of Glu924 and a ribose hydroxyl group in the RNA present in the nucleotide-free structure of WT MDA5 is lost in the I923V nucleotide-free structure. The loss of this protein-RNA contact partly explains the reduced RNA binding affinity and protein-RNA interaction area of the I923V variant reported above.

**Fig. 4.**
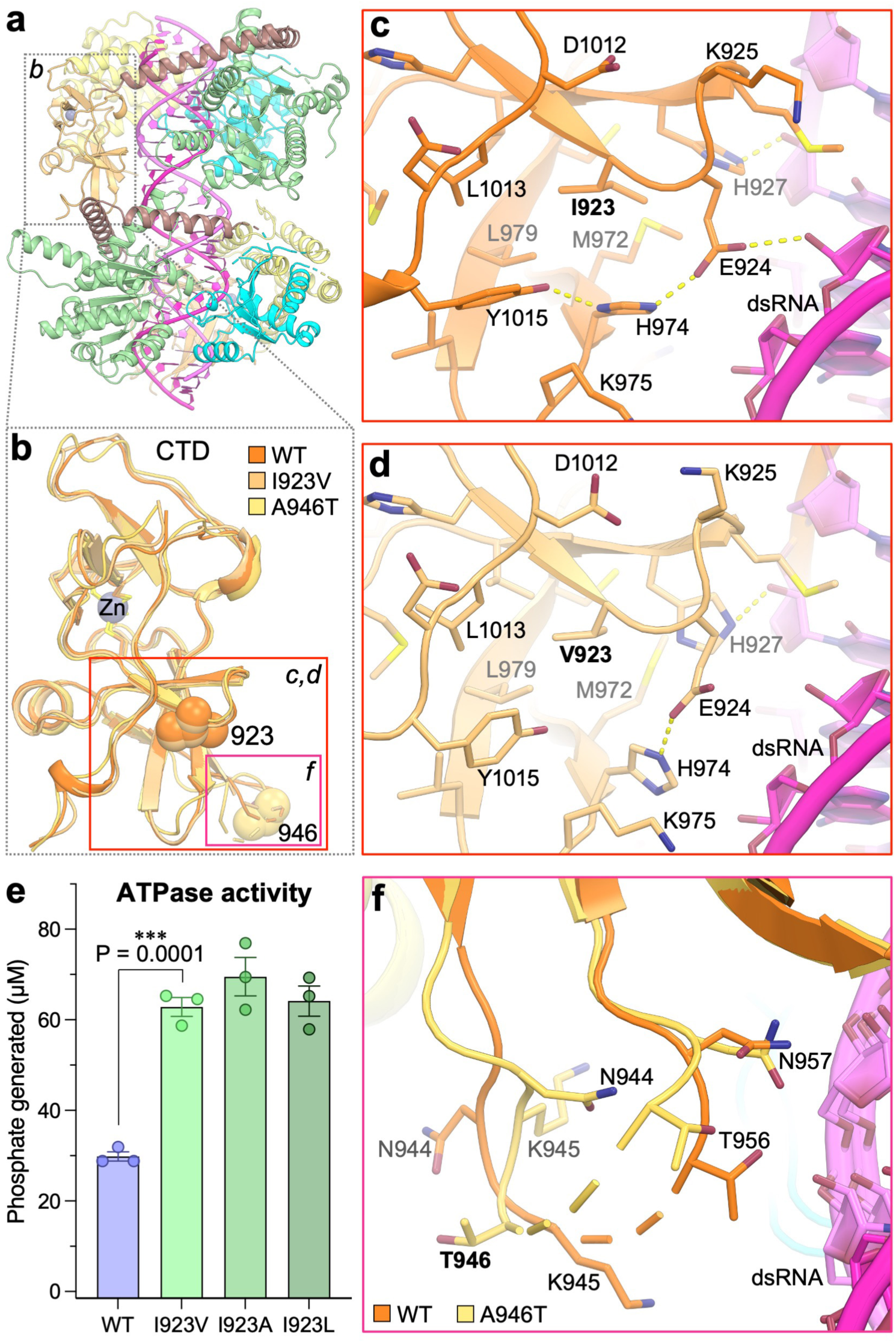
Isoleucine is required at position 923 to sterically regulate ATPase activity. (**A**) Overall structure of the MDA5-dsRNA filament without nucleotide. (**B**) The superimposed C-terminal domains of WT, I923V, and A946T MDA5 from the cryo-EM structures without nucleotide (PDB IDs 6H61, 9F0J, and 9F3P). (**C**) Closeup of Ile923 and surrounding residues in the WT MDA5 structure. (**D**) Closeup of Val923 and surrounding residues in the MDA5 I923V structure. (**E**) ATPase activities of WT, I923V, I923A, and I923L MDA5. (**F**) Closeup of the loop containing residue 946 in the superimposed structures of WT and A946T MDA5.

Considering the 3-fold increase in ATPase activity associated with relatively subtle structural changes in the I923V variant, we measured the ATPase activities of MDA5 proteins with slightly more or less conservative mutations at position 923, substituting either alanine or leucine for isoleucine. We found that the I923A and I923L mutants had the same ATPase activity as I923V (**Figure 4E**). Hence, further decreasing the bulk of the hydrophobic side chain from valine to alanine did not further increase ATPase activity, and substitution with leucine, an isomer of the isoleucine residue found in WT MDA5, did not reduce ATPase to the level of WT MDA5. We conclude that the isoleucine side chain at position 923 regulates the conformational changes necessary for ATP hydrolysis by shaping the CTD fold to tune ATPase activity to an optimal evolved level in WT MDA5.

In contrast to residue 923, residue 946 is located within a partly disordered solvent exposed loop that does not contribute to the core fold of the CTD. This loop is poorly defined in the cryo-EM maps. There are minor differences in the conformation of the loop in the WT and A946T structures, but these differences do not alter the protein fold, RNA binding interface or active site (**Figure 4F**). This is consistent with the absence any significant effects of the A946T substitution on the overall structure, filament assembly, and ATPase activity of MDA5. Together, our data suggest that the A946T substitution has no direct effect on the structural or biochemical properties of MDA5. We note that threonine is a phosphorylatable residue raising the possibility that T1D protection associated with the T946A allele may be due to a difference in phosphorylation state.

## Discussion

We have shown here that T1D-protective MDA5 variants have pleiotropic effects on the structural and biochemical activities of MDA5 (**Figure 5**). The rare variants E627* and I923V reduce the MDA5-dependent cellular IFN-β response to picornavirus infection, in agreement with previous studies, but we found that variants R843H and T946A, in combination, as they are predominantly found in humans due to linkage disequilibrium, had no significant effect on IFN-β expression. For the E627* variant, the deletion of most of the helicase module of MDA5 is sufficient to explain the loss of RNA binding, destabilization of the protein fold, and therefore the loss of signaling function. In contrast, we find that the I923V variant has increased ATPase activity. Our cryo-EM structures of the I923V variant bound to dsRNA at different stages of ATP hydrolysis reveal that the Ile923 sidechain regulates the conformational changes necessary for ATP hydrolysis. The increased ATPase activity of the I923V variant may be attributed to the slight reduction of the surface complementarity of the MDA5 RNA binding interface in I923V variant. As a result, the I923V variant has altered helical twist distributions of MDA5-dsRNA filaments in the ATP-bound and ADP-AlF_4_-bound states. Since engineered I923A and I923L variants had the same hyperactive ATPase activity as I923V, we propose that the ancestral isoleucine side chain at position 923 functions as a molecular brake, regulating the conformational changes necessary for ATP hydrolysis by shaping the CTD fold to tune ATPase activity to an optimal evolved level in WT MDA5. The increased ATPase activity of the I923V variant will result in overzealous proofreading, promoting premature dissociation from dsRNA. This in turn restricts the formation of active signaling complexes on dsRNA.

**Fig. 5.**
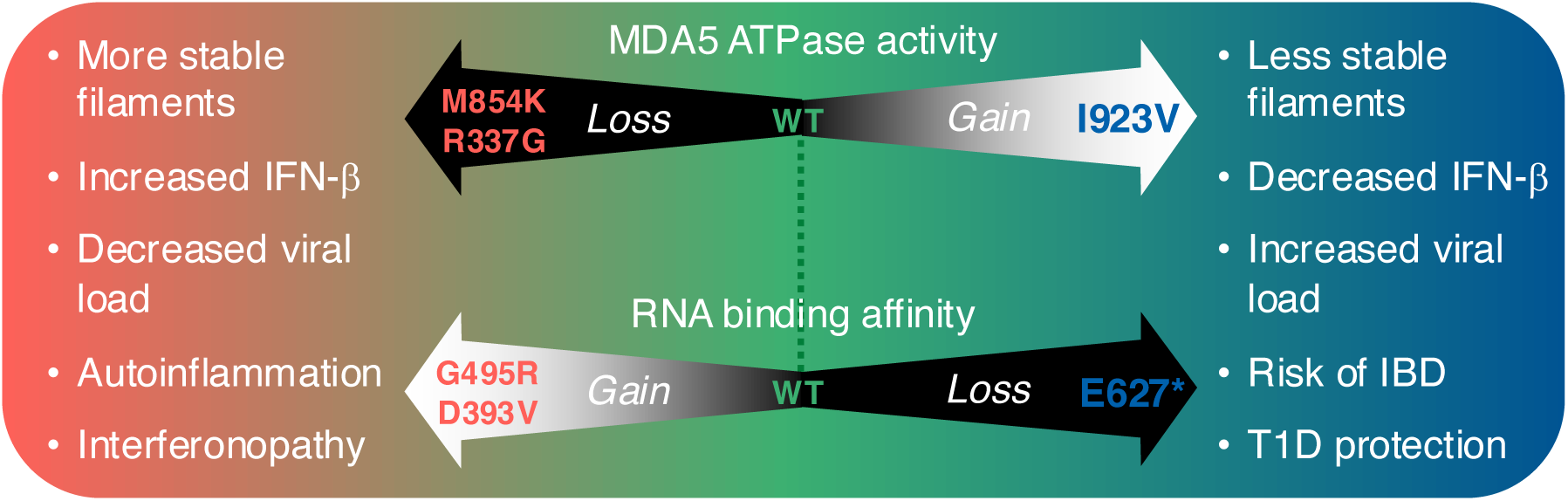
MDA5 variants tune immune homeostasis via a fitness trade-off between viral clearance and tissue damage. T1D-protective MDA5 variants E627* and I923V lead to loss of signaling function. E627* causes loss of RNA binding whereas I923V hyperactivates ATPase activity, which promotes dissociation from RNA. Loss of signaling function is associated with increased viral loads and IBD onset, while reducing autoimmune tissue damage. Variants that increase RNA binding or inhibit ATPase activity, leading to a gain of signaling function and autoinflammation, but infections are cleared more effectively.

The R843H and A946T substitutions had no apparent effects on ATPase activity, structure, dsRNA binding affinity, or filament formation on dsRNA in our assays, leaving the molecular basis of T1D-protective effects of these mutants unclear. A new meta-analysis of recent, large GWAS datasets confirmed association of E627*, I923V, T946A, and R843H with significant levels of protection against T1D as well as other autoimmune-related diseases including psoriasis and hypothyroidism (Wallace *et al*, 2024). Fine-mapping suggested that the effect associated with the R843H variant was not independent and was likely a result of linkage with the T946A variant (Wallace *et al*., 2024), consistent with previous studies concluding that associations with R843H could be explained by its co-occurrence with T946A (Gorman *et al*., 2017; Nejentsev *et al*., 2009). Together, the available evidence and analyses suggest that R843H is not independently protective against T1D and that T946A is protective via an indirect mechanism. We hypothesize that a threonine residue at position 946 can be phosphorylated or otherwise post-translationally modified to promote signaling. For example, a phosphothreonine at position 946 may contribute to the recruitment of signaling cofactors (e.g. PACT, ZCCHC3, TRIM65, or K63-linked ubiquitin chains), or alternatively increase the lifetime of MDA5 in the cytosol (e.g. by protecting it from degradation).

In conclusion, the T1D-protective MDA5 variants E627*, I923V, and T946A each ultimately lead to a loss of MDA5-dependent signaling but do so via three distinct mechanisms. The net loss-of-function effect that T1D-protective mutations have on interferon signaling is the converse of MDA5 variants associated with autoinflammatory disease, which have a gain-of-function effect on signaling and also act via multiple distinct mechanisms (Rice *et al*., 2014; Yu *et al*., 2021). Notably, in addition to protecting against autoimmune-related diseases, these loss-of-function variants increased the risk of viral infection and inflammatory bowel disease (IBD), specifically Crohn’s disease and ulcerative colitis (Wallace *et al*., 2024). Furthermore, the degree of T1D protection and IBD risk were closely correlated, suggesting that MDA5 loss-of-function variants offer a fundamental fitness trade-off between viral clearance and tissue damage (Wallace *et al*., 2024). Robust clinical links have emerged between enteric virus infection and the onset of both T1D (Hyoty & Taylor, 2002; Yeung *et al*, 2011) and IBD (Axelrad *et al*, 2019). Indeed, MDA5-induced inflammation and cell death in the pancreas following rotavirus infection contributes to autoimmune destruction of pancreatic β-cells (Dou *et al*., 2017), and gastrointestinal infection has been associated with increased risk of developing IBD in a large clinical study (Axelrad *et al*., 2019). We propose a model in which loss-of-function MDA5 variants protect against T1D by reducing autoimmune β-cell killing triggered by MDA5-dependent IFN-β production and inflammation following viral infection, while also contributing to the induction of IBD (Axelrad *et al*., 2019; Wallace *et al*., 2024) due to increased susceptibility to viral infection from the loss of MDA5 antiviral activity.

## Materials and Methods

### RNA synthesis

RNAs were transcribed in vitro using the MEGAscript T7 Transcription Kit (Invitrogen, cat. no. AM1333) or HiScribe T7 High Yield RNA Synthesis Kit (New England BioLabs, cat. no. E2040S) following the manufacturers’ protocols. The 200-bp, 300-bp, and 1-kb dsRNA contained the first 200, 300 or 1000 bases of the mouse IFIH1 gene, respectively. The complementary RNA strands were transcribed from DNA templates with a preceding 5′ TAATACGACTCACTATAG 3′ sequence and a T7 promoter on the coding strand. The in vitro transcription reactions were performed at 37°C for 2, 4, 6 h or overnight. Transcripts were treated with Turbo DNase and purified with the PureLink RNA Mini Kit (ThermoFisher, cat. no. 12183018A) or the Monarch RNA Cleanup Kit (500 μg) (New England BioLabs, cat. no. T2050L). Samples were eluted in nuclease-free duplex annealing buffer (30 mM HEPES pH 7.5, 100 mM KCl (Integrated DNA Technologies)). Eluted transcripts were incubated at 95°C for 5 min and cooled to room temperature over 2 h to eliminate secondary structure and enable annealing of complementary strands of RNA.

### Cell lines expressing MDA5 T1D-protective variants

RIG-I KO A549 cells expressing ACE2 (A549 RIG-I^−/−^ ACE2^+^ cells) were transduced with a recombinant lentivirus to express T1D-protective MDA5 variants in a doxycycline-dependent manner. To generate the lentiviruses, HEK293T cells grown in Dulbecco’s modified Eagle’s medium (DMEM; Lonza), high glucose media with GlutaMAX Supplement (Gibco) were transfected with a pLVX-TetOne-Puro vector (Takara) containing a gene encoding human MDA5 (UniProt: Q9BYX4) with an N-terminal Flag tag. HEK293T cells were transfected (24 h after seeding 4-5 x10^6^ cells in 8 ml of medium in a 10-cm plate) with 7.0 µg of pLVX-TetOne-Puro-hMDA5 vector in 600 µl sterile water with Lenti-X Packaging Single Shots (Takara). 16 h post-transfection, 6 ml of fresh complete growth medium was added. After a further 48 h incubation, cells were harvested by centrifuging at 500 g for 10 min. Supernatant containing viral particles was filtered through a 0.45-μm filter and the lentivirus titer determined by enzyme-linked immunosorbent assay (ELISA) with the Lenti-X p24 Rapid Titer (Single Wash) Kit (Takara). To generate cell lines expressing MDA5 variants, A549 RIG-I^−/−^ ACE2^+^ cells (80-90% confluent in 6-well plates) were transduced by adding 10 µg ml^−1^ Polybrene (Tocris) and recombinant lentivirus to a multiplicity of infection (MOI) between 2 and 10, followed by centrifugation at 800 g for 30 min. After 16 h the transduction medium was exchanged for fresh growth medium. After a further 48 h incubation, 2 µg ml^−1^ puromycin was added and antibiotic selection maintained for two weeks. MDA5 expression was assessed in the presence of 0-100 µg ml^−1^ doxycycline by Western blotting with anti-FLAG (Sigma-Aldrich, RRID:AB_262044, 1:5000 dilution) or anti-MDA5 antibody (Enzo Life Sciences, RRID:AB_2893162, 1:1000 dilution).

### Virus infection assays

Recombinant encephalomyocarditis virus (EMCV, strain Mengovirus) Leader-Zn mutant (carrying mutations C19A and C22A in the zinc finger domain of the viral Leader protein (Hato *et al*., 2007)) was generated by transfecting RNA produced from an infectious clone into BHK21 cells. Viruses were harvested after complete cytopathogenic effect, concentrated by ultracentrifugation (30% sucrose, 140,000 g for 16 h, 4°C, SW32Ti rotor), diluted in PBS and stored at −80°C. Virus titers were determined by end point titration according to the method of Spearman-Kärber and expressed as 50% Tissue Culture Infectious dose (TCID50). EMCV was used to infect A549 RIG-I^−/−^ ACE2^+^ cells expressing MDA5 variants cultured in DMEM containing sodium pyruvate and Glutamax (Gibco, cat. no. 2206106), supplemented with 1 μg ml^−1^ doxycycline, 10% fetal bovine serum (FBS), and 1% Pen-Strep (Lonza). Concentrations for *IFNB1* mRNA, *IFIT1* mRNA, and EMCV viral RNA (vRNA) were quantified by RT-qPCR from total RNA extracted from the infected cells 7 h post infection as described previously (Bruurs *et al*, 2023) (**Figure S1**).

### MDA5 protein purification

A gene encoding mouse MDA5 (Ifih1, UniProt: Q8R5F7) was cloned into the pET28a vector with an N-terminal hexa-histidine tag followed by a tobacco etch virus (TEV) protease cleavage site as described (Berke & Modis, 2012). MDA5 residues 646-663, in the flexible L2 surface loop of the helicase 2 insert domain (Hel2i), were deleted for solubility, resulting in a 114-kDa polypeptide chain. The ΔL2 loop deletion does not affect the dsRNA binding, ATPase or interferon signaling activities of MDA5 (Berke & Modis, 2012; Wu *et al*., 2013; Yu *et al*., 2018). The T1D-protective mutations were introduced into the MDA5-ΔL2 construct with the Q5 Site-Directed Mutagenesis Kit (New England BioLabs).

Escherichia coli Rosetta(DE3)pLysS cells (Novagen, cat. no. 71403) were transformed with an MDA5 construct and grown in 2xTY medium to OD_600_ 0.4-0.6 at 37°. After cooling to 16°C, protein expression was induced with 0.5 mM isopropyl-β-D-1-thiogalactopyranoside (IPTG) overnight at 16°C. Harvested cells were resuspended in lysis buffer (30 mM HEPES pH 7.7, 150 mM M NaCl, 5% glycerol, 1 mM Tris(2-carboxyethyl)phosphine (TCEP), with cOmplete EDTA-free Protease Inhibitor Cocktail (Roche, cat. no. 11873580001) and 1 U ml^−1^ Benzonase Nuclease (Merck, cat. no. 70746). Cells were lysed by sonication (5s s on, 10 s off, 40% power), and the lysate was centrifuged at 37,500 g for 1 h. The supernatant was loaded onto a pre-equilibrated 5 ml HisTrap HP column (Cytiva, cat. no. 17-5248-02), washed with wash buffer (30 mM HEPES pH 7.7, 500 mM NaCl, 20 mM imidazole, 5% glycerol, and 1 mM TCEP). MDA5 was eluted with Ni-NTA elution buffer (20 mM HEPES 7.7, 150 mM NaCl, 300 mM imidazole, 5% glycerol, 1 mM TCEP). MDA5 was further purified on a Resource Q anion exchange column (Cytiva) (buffer A: 20 mM HEPES 7.7, 50 mM NaCl, 1 mM dithiothreitol (DTT); buffer B: 20 mM HEPES 7.7, 1 M NaCl, 1 mM DTT), and a Superdex 200 Increase 10/300 GL size-exclusion column (Cytiva) in SEC buffer (20 mM HEPES pH 7.7, 150 mM M KCl, 1 mM DTT, 5% glycerol). Purified protein was used immediately for cryo-EM grid preparation and ATPase assays, or flash-frozen and stored at −80°C.

### Nanoscale Differential Scanning Fluorimetry (nanoDSF)

1 µM MDA5 in SEC buffer (20 mM HEPES pH 7.7, 150 mM KCl, 1 mM DTT), alone or incubated with 15 ng µl^−1^ (22 nM) 1-kb dsRNA for 30 min at room temperature, was loaded into standard capillaries (NanoTemper, #PR-C002). Intrinsic protein fluorescence at 330 nm and 350 nm, F330 and F350, respectively, was measured from 15°C to 80°C, with a heating rate of 1°C per minute, with a Prometheus NT.48 nano-fluorimeter (NanoTemper). The melting temperatures (T_m_ values) were calculated from changes in the fluorescence ration (F350/F330) using the PR.Stability Analysis v1.1 software (NanoTemper).

### Bio-layer interferometry (BLI)

BLI experiments were performed on an Octet Red384 (ForteBio Inc.) instrument. Binding experiments were carried out at 30°C in assay buffer (20 mM HEPES pH 7.7, 150 mM KCl and 1 mM DTT, 2 mg ml^−1^ BSA). BSA was required in the assay buffer to avoid non-specific binding to the sensor.

Pierce RNA 3’ End Biotinylation Kit (ThermoFisher, cat. no. 20160) was used to label the 3’ end of 200-bp, 300-bp or 1-kb RNA duplexes (produced as described above) with biotin. Biotinylated RNA (2.5 μg ml^−1^) was immobilized onto Octet streptavidin (SA) Biosensors (Sartorius), and binding performed for varying concentrations of MDA5 (**Figure S2**). Sensors were hydrated in assay buffer for at least 600 s prior to all measurements. Binding experiments comprised sensor equilibration (60 s), loading (600 s), baseline (90 s), and association and dissociation (600 s each) steps. Data analysis was performed using the Octet Analysis Studio v 11.1 software (Sartorius). Loaded sensors dipped into assay buffer during the association and dissociation steps were used as references and subtracted from all samples during analysis to correct for baseline drift. Local kinetic fitting was used to determine K_d_ values from binding curves with 125 nM MDA5. K_d_ values obtained from three or four replicate runs were averaged, and standard error calculated, using GraphPad Prism v10.

### ATPase assay

ATPase activities were measured using the ATPase/GTPase Activity Assay Kit (Sigma-Aldrich, cat. no. MAK113). Reactions containing 180 nM MDA5 and 25 ng µl^−1^ 1-kb RNA in buffer (20 mM HEPES pH 7.7, 150 mM KCl, 4 mM ATP, 4 mM MgSO_4_, 1 mM DTT) were incubated at 37°C for 15 min and quenched by the addition of malachite green. Reactions were performed in clear, flat-bottom 96-well plates, and the inorganic phosphate produced by ATP hydrolysis was monitored by tracking absorbance at 620 nm using a CLARIOstar microplate reader (BMG Labtech). Results were analyzed with Prism v9.5.1 (GraphPad, graphpad.com), and for mutants, data was normalized relative to the phosphate generated by wild type MDA5.

### Negative stain EM

200 nM MDA5, 3 ng µl^−1^ (4.4 nM) 1-kb dsRNA, and 4 mM ATP were incubated at room temperature for 30 min in 20 mM HEPES pH 7.7, 150 mM KCl, 1 mM DTT, 5% glycerol. Carbon film 300-mesh grids (Agar Scientific) were glow discharged at 25 mA for 1 min. Samples were applied to the grids, washed with RNase-free water, stained using uranyl acetate [2% (w/v)], and imaged with a 120 kV Technai G2 Spirit electron microscope (ThermoFisher). Images were taken at −2 to −4 µm defocus and 26,000× magnification (4 Å pixel^−1^).

### Cryo-EM sample preparation and data collection

1 g l^−1^ purified MDA5 protein was incubated with 0.05 g l^−1^ 1-kb dsRNA in 20 mM HEPES pH 7.7, 100 mM KCl, 5 mM MgCl_2_, 2 mM DTT, and nucleotide (10 mM ATP; 10 mM AMPPNP; or 2 mM ADP, 4 mM AlCl_3_ and 40 mM NaF) on ice for 2-3 min. Samples were diluted twofold with the same buffer and 3.5 µl of sample was immediately applied onto a glow-discharged 300-mesh gold Quantifoil R1.2/1.3 grid (Quantifoil Micro Tools). Grids were glow discharged with an Edwards 12E6/531 glow discharger at 30 mA for 60 s. Grids were blotted for 2-4 s, held for a 15 s wait time, and plunge-frozen in liquid ethane cooled by liquid nitrogen with a Vitrobot Mark IV (ThermoFisher) at 4°C and 100% humidity.

Cryo-EM data were collected on 300 kV Titan Krios microscopes (ThermoFisher) equipped with Gatan K3 detectors and Gatan BioQuantum energy filters at the MRC Laboratory of Molecular Biology and EMBL Heidelberg. Movies were recorded with a fluence of 40-48 electrons per square angstrom (e^−^ Å^−2^), an average exposure of 1.0 e^−^ Å^−2^ per frame, and a flux of 5-5.7 e^−^pixel^−1^ s^−1^. A 20-eV energy selection slit width was used. The nominal defocus value ranged from −0.5 to −2.5 µm in 0.5 µm increments. All I923V ATP samples were collected on a K3 detector (Gatan) at 105,000x magnification (0.826 Å pixel^−1^). The A946T sample was collected at 96,000x magnification (0.822 Å pixel^−1^). Detectors were used in counting mode. Data were acquired with EPU and two shots per hole, except for the A946T AMPPNP sample, which was acquired with SerialEM and six shots per hole. See **Table S1** for additional data collection parameters.

### Image processing and helical reconstruction

Movies were motion-corrected and dose-weighted with MotionCor2.0 in Relion4.0 (Kimanius *et al*, 2021). The contrast transfer function was estimated with CtfFind4.1 (Rohou & Grigorieff, 2015) and the micrographs were aligned without dose weighting. Image reconstruction with helical symmetry averaging was performed in Relion4.0 (He & Scheres, 2017). Segments were picked with crYOLO (Wagner *et al*, 2019) from a template trained on the M854K ATP dataset (EMD-12213) (Yu *et al*., 2021). Segment were subjected to several rounds of 2D and 3D classification. 3D refinement, CTF refinement, Bayesian polishing, and post-processing were performed in Relion4.0 (Kimanius *et al*., 2021; Scheres, 2014). To calculate the helical twist distribution of helical segments, three independent rounds of 3D classification were performed with five classes. Segments were then placed in Low twist (71°–81°), Intermediate twist (81°– 91°) or High twist (91°–96°) bins for plotting histograms of the twist distribution with Prism v9.5.1 (GraphPad). The number of segments contributing to each 3D class was weighted evenly. The numbers of segments used in 3D classification for twist distribution calculation datasets were as follows: I923V ATP dataset, 457,811 segments; I923V AMPPNP dataset, 269,092 segments; I923V no-nucleotide dataset, 547,562 segments; and I923V ADP-AlF_4_ dataset, 683,412 segments. See **Table S1** for the initial and final number of segments used for each dataset and for the final resolution and helical parameters of each reconstruction (**Figure S3**).

### Model building and refinement

Previously determined cryo-EM structures of MDA5-dsRNA filaments with similar helical symmetry (PDB:7BKP, [http://doi.org/10.2210/pdb7BKP/pdb], PDB7NIC, [http://doi.org/10.2210/pdb7NIC/pdb], or PDB:7NIQ, [http://doi.org/10.2210/pdb7NIQ/pdb]) (Yu *et al*., 2018) were used as the starting atomic models for model building. The model was docked into the density for the central subunit in each map with the Fit in Map function of UCSF Chimera (Pettersen *et al*, 2004). The docked models were rebuilt with COOT (Emsley & Cowtan, 2004). Models of adjacent protomers were generated in Chimera by applying the helical symmetry calculated in Relion. The resulting models with three MDA5 subunits were used for subsequent real space refinement in Phenix 1.21 (Liebschner *et al*, 2019). Real space refinement in Phenix included global minimization and atomic displacement parameter refinement, incorporating restraints on secondary structure, sidechain rotamers, mainchain torsion angles, and non-crystallographic symmetry between the three modeled protein subunits (Liebschner *et al*., 2019).

To determine which conformational state the helicase modules were in, each model was superimposed onto the fully closed structure of LGP2 (PDB:5JAJ (Uchikawa *et al*., 2016)) using the secondary structure elements of Hel1 as the reference. The conformational state of the helicase domain was defined based on the rotation angles relating the Hel2 domains of the aligned structures as follows: closed state, <5° angle; semi-closed state, >5° angle. Protein interfaces were analyzed with the Protein interfaces, surfaces, and assemblies (PISA) service at the European Bioinformatics Institute [http://www.ebi.ac.uk/pdbe/prot_int/pistart.html] (Krissinel & Henrick, 2007).

### Statistical analysis

No statistical methods were used to predetermine sample size, experiments were not randomized, and the investigators were not blinded to experimental outcomes. Cell signaling data and BLI data are represented as the mean ± standard error of the mean of three replicates conducted in three independent experiments. ATPase assays were performed at least three times in independent experiments. Scatter plots, histograms and error bars were plotted with GraphPad Prism 10.2.3 and Microsoft Excel v.16.84. Statistical significance was assessed using unpaired two-tailed t-tests assuming Gaussian distributions (without Welch’s correction). Statistical significance was assigned as follows: n.s., P > 0.05; *, P < 0.05; **, P < 0.01; ***, P < 0.001.

## Data availability

The atomic coordinates were deposited in the Protein Data Bank with accession codes 9F2W at https://doi.org/10.2210/pdb9f2w/pdb, 9F2L https://doi.org/10.2210/pdb9f2l/pdb, 9F1U https://doi.org/10.2210/pdb9F1U/pdb, 9F20 https://doi.org/10.2210/pdb9f20/pdb, 9F0J https://doi.org/10.2210/pdb9f0j/pdb, and 9F3P https://doi.org/10.2210/pdb9f3p/pdb. The cryo-EM densities were deposited in the EM Data Bank with codes EMD-50165, EMD-50150, EMD-50136, EMD-50137, EMD-50111, and EMD-50175. Other data underlying this article are available in the article, Supplementary Figures and Table, and online supplementary data.

## Acknowledgments

We thank the following facility staff for assistance in cryo-EM data collection: Bilal Ahsan, Giuseppe Cannone, Shaoxia Chen, Grigory Sharov, and other staff at the MRC-LMB EM Facility; and Felix Weis (EMBL Heidelberg Cryo-EM Service Platform). We thank Stephen McLaughlin and Chris Batters (MRC-LMB Biophysics Facility) for assistance with Octet experiments. We thank Tetsuo Hasegawa and members of the Modis lab for insightful discussions. We thank MRC-LMB Scientific Computing for computing support. We acknowledge the support of the MRC-LMB Media & Glass Wash facility. This work was supported by the Wellcome Trust [101908/Z/13/Z to Y.M.; 217191/Z/19/Z to Y.M.; 215378/Z/19/Z to R.S.], and the Human Frontier Science Program [LT000454/2021-L to A.H.d.V.). Open access publication was funded by the University of Cambridge.

## Author Contributions

Conceptualization: R.S., A.d.H.V., B.J.F., Y.M. Formal Analysis: R.S., A.d.H.V., J.D.J., M.Z., F.J.M.v.K., Y.M. Methodology: R.S., A.d.H.V., J.D.J., M.Z., B.J.F., F.J.M.v.K., Y.M. Investigation: R.S., A.d.H.V., J.D.J., M.Z., Y.M. Visualization: R.S., Y.M. Funding acquisition: R.S., A.d.H.V., Y.M. Project administration: Y.M. Supervision: B.J.F., F.J.M.v.K., Y.M. Writing—original draft: Y.M. Writing—review & editing: R.S., A.d.H.V., B.J.F., F.J.M.v.K., Y.M.

## Competing Interest Statement

Y.M. is a consultant for Related Sciences LLC. None of the other authors have any conflicts of interest.

## Notes

https://doi.org/10.2210/pdb9f2w/pdb

https://doi.org/10.2210/pdb9f2l/pdb

https://doi.org/10.2210/pdb9F1U/pdb

https://doi.org/10.2210/pdb9f20/pdb

https://doi.org/10.2210/pdb9f0j/pdb

https://doi.org/10.2210/pdb9f3p/pdb

