## Supplementary Figures and Table (Table S1 and Figs S1-S3) for "Molecular basis of autoimmune disease protection by MDA5 variants"

**Table S1.** Cryo-EM data collection and structure determination parameters.

| | **I923V No nt.** | **I923V ATP** | **I923V ADP-$AlF_4$** | | | **A946T No nt.** |
|---|---|---|---|---|---|---|
| **Data Collection and Processing** | | | | | | |
| Microscope/Detector | Krios/K3 | Krios/K3 | | Krios/K3 | | Krios/K3 |
| Micrograph fluence ($e^- Å^{-2}$) | 40 – 48 | 40 – 48 | | 40 – 48 | | 40 – 48 |
| Exposure per frame ($e^- Å^{-2}$) | 1.0 | 1.0 | | 1.0 | | 1.0 |
| Nominal defocus range (µm) | -0.5 – -2.5 | -0.5 – -2.5 | | -0.5 – -2.5 | | -0.5 – -2.5 |
| Pixel size (Å) | 0.826 | 0.826 | | 0.826 | | 0.822 |
| N. initial segment images | 1,008,651 | 683,122 | | 1,406,812 | | 1,378,603 |
| **Map averaging and refinement** | | | **73˚ twist** | **81˚ twist** | **88˚ twist** | |
| N. final segment images | 547,562 | 337,327 | 53,484 | 347,343 | 98,676 | 234,719 |
| Resolution, unmasked ½-maps (Å) | 3.57 | 4.35 | 4.25 | 4.05 | 4.48 | 4.25 |
| Final resolution with masking (Å) | 3.33 | 4.21 | 3.86 | 3.67 | 4.13 | 3.97 |
| Max. local resolution range (Å) | 5.94-3.12 | 7.31-3.80 | 7.91-3.48 | 6.58-3.35 | 7.85-3.84 | 6.52-3.67 |
| Map sharpening B factor ($Å^2$) | -90 | -130 | -50 | -85 | -90 | -150 |
| Helical twist (˚) | 91.63 | 84.15 | 73.94 | 80.71 | 88.20 | 89.44 |
| Helical rise (Å) | 44.84 | 43.92 | 42.78 | 43.86 | 44.50 | 44.58 |
| **Atomic model fit in data** | | | | | | |
| CC (mask), Phenix v1.21 | 0.81 | 0.74 | 0.78 | 0.73 | 0.74 | 0.76 |
| CC (volume), Phenix v1.21 | 0.80 | 0.74 | 0.78 | 0.73 | 0.73 | 0.75 |
| **Atomic model composition** | | | | | | |
| N. non-hydrogen atoms | 5782 | 5,981 | 6,129 | 5,979 | 6,128 | 6,067 |
| Protein residues | 640 | 675 | 686 | 673 | 681 | 682 |
| RNA nucleotides | 30 | 28 | 28 | 28 | 30 | 30 |
| Ligand | None | ATP | ADP-$AlF_4$ | ADP-$AlF_4$ | ADP-$AlF_4$ | None |
| $Zn^{2+}$ ions | 1 | 1 | 1 | 1 | 1 | 1 |
| **Atomic model geometry, ADPs** | | | | | | |
| Bond lengths (Å) | 0.004 | 0.003 | 0.003 | 0.004 | 0.003 | 0.003 |
| Bond angles (˚) | 0.525 | 0.566 | 0.578 | 0.606 | 0.656 | 0.548 |
| Protein min/max/mean ADP | 26/150/94 | 76/196/140 | 82/163/117 | 80/201/147 | 90/218/160 | 41/163/106 |
| Nucleotide min/max/mean ADP | 17/72/41 | 57/90/71 | 67/79/72 | 63/106/83 | 88/119/100 | 32/62/41 |
| Ligand min/max/mean ADP | None | 136/224/138 | 122/188/126 | 147/221/151 | 151/215/157 | None |
| **Validation** | | | | | | |
| MolProbity score, Phenix v1.21 | 1.82 | 2.00 | 1.70 | 2.03 | 2.11 | 2.06 |
| Clash score, Phenix v1.21 | 8.53 | 11.78 | 6.71 | 13.30 | 14.96 | 11.67 |
| Rotamer outliers (%) | 0.00 | 0.00 | 0.00 | 0.00 | 0.00 | 0.00 |
| Ramachandran plot | | | | | | |
| % favored | 94.9 | 93.8 | 95.3 | 94.1 | 93.5 | 92.3 |
| % allowed | 5.1 | 6.2 | 4.7 | 5.9 | 6.5 | 7.7 |
| % outliers | 0.0 | 0 | 0.0 | 0.0 | 0.0 | 0.0 |
| PDB code | 9F0J | 9F2W | 9F2L | 9F1U | 9F20 | 9F3P |
| EMDB code | EMD-50111 | EMD-50165 | EMD-50150 | EMD-50136 | EMD-50137 | EMD-50175 |

*ADPs, atomic displacement parameters

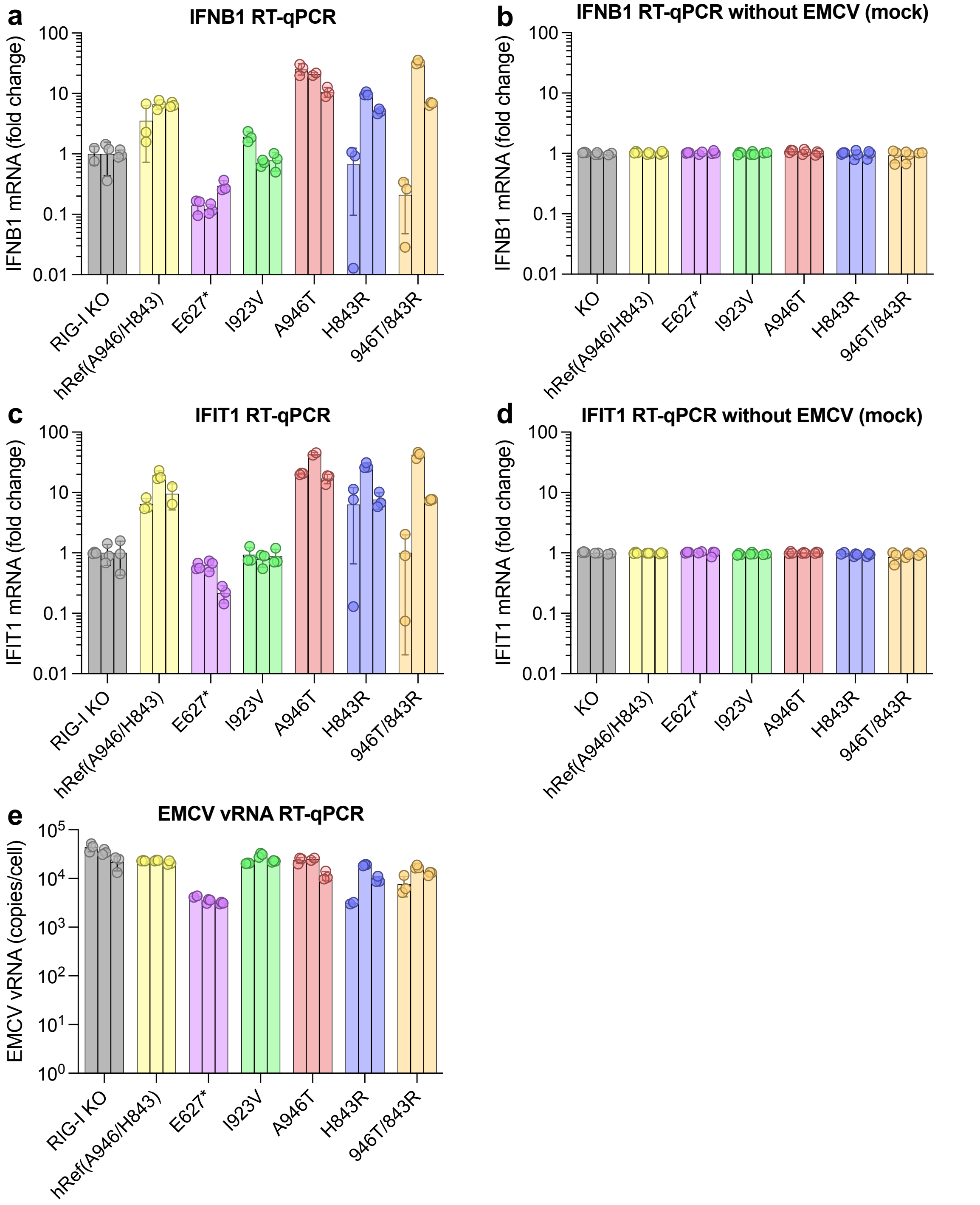


**Fig. S1.** EMCV data with technical replicates shown and EMCV mock infections. (**A,B**) RT-qPCR quantification of IFNB1 mRNA in A549 RIG-I KO cells stably expressing the indicated MDA5 variant under a doxycycline-inducible promoter 7 h after infection with encephalomyocarditis virus (EMCV), (**A**), or 7 h after addition of mock buffer, (**B**). hRef, human reference sequence. (**C,D**) RT-qPCR quantification of IFIT1 mRNA after EMCV infection, (**C**), or without EMCV infection, (**D**). (**E**) RT-qPCR quantification of EMCV RNA. All technical replicates from three independent qPCR experiments are shown. See **Data S1** for source data.

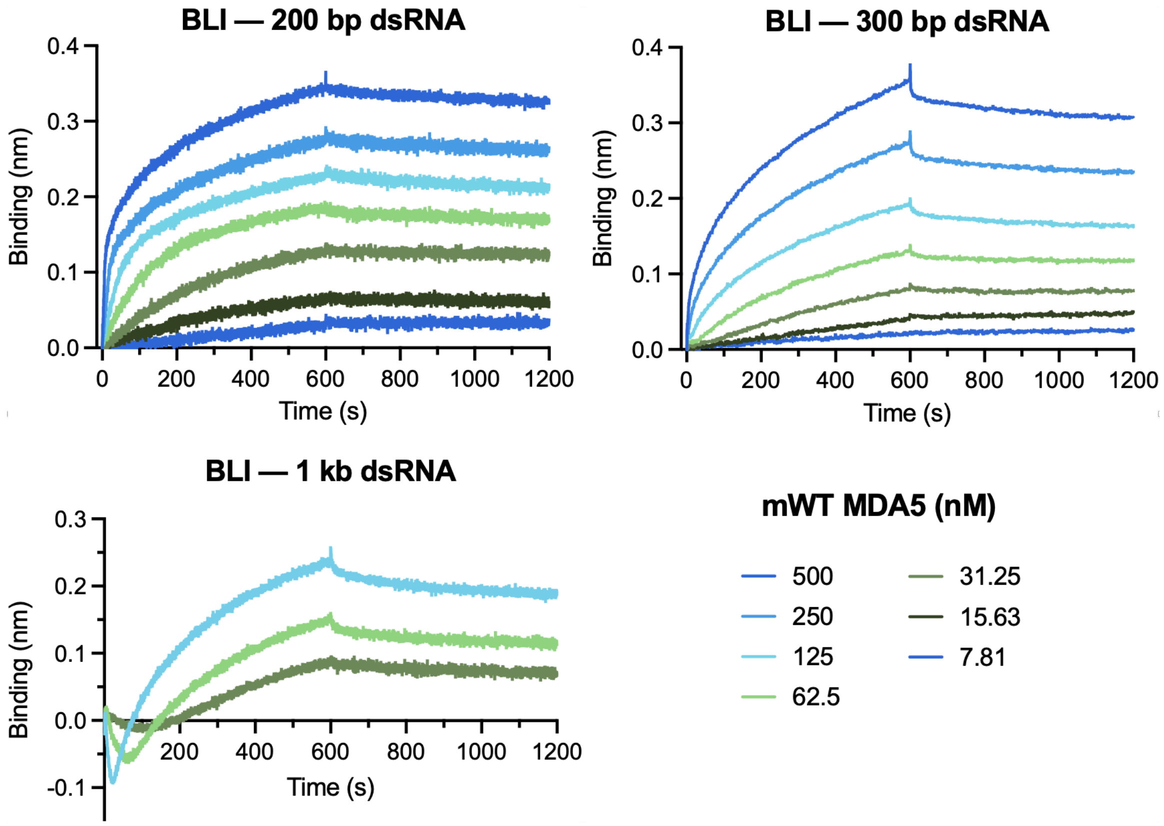


**Fig. S2.** Bio-layer interferometry (BLI) with different concentrations of mouse wild type (mWT) MDA5. 3'-biotinylated dsRNA was immobilized on a streptavidin sensor and MDA5 was added to the mobile phase. The curves for 1-kb dsRNA have a complex shape in the first 100 s of the binding phase.

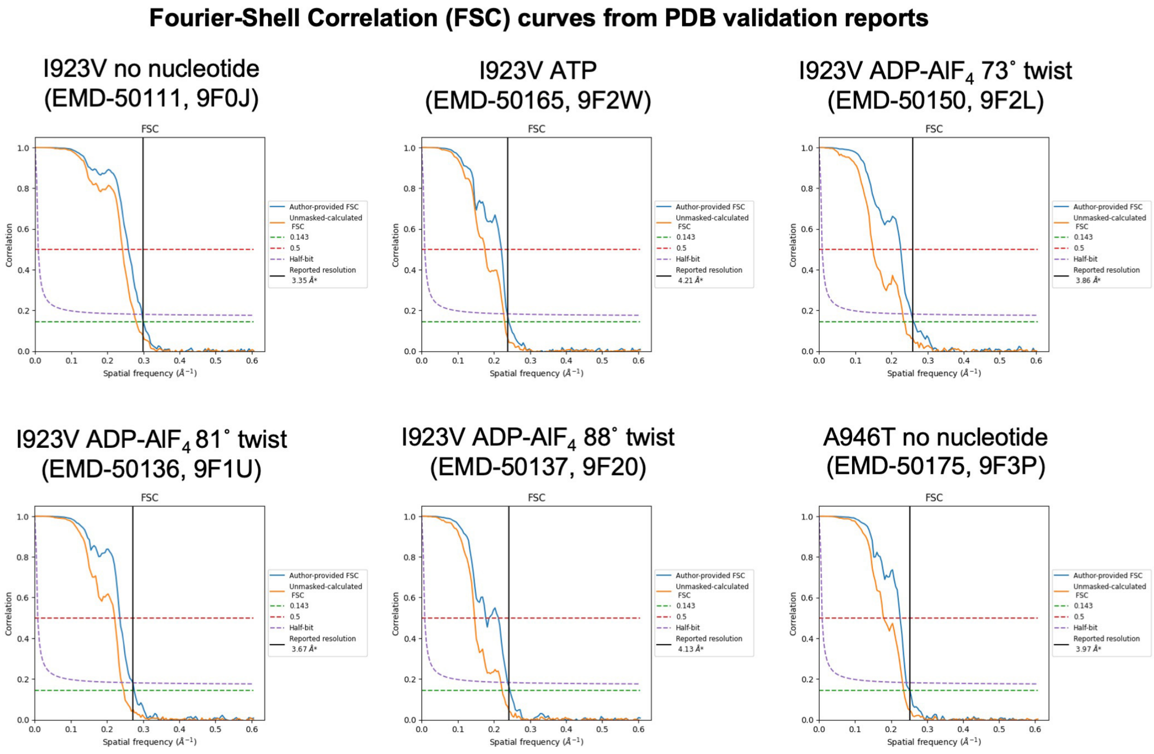


**Fig. S3.** Fourier shell correlation curves for cryo-EM image reconstructions. Electron Microscopy Data Bank (EMDB) and Protein Data Bank (PDB) accession codes are listed for each structure. Graphs were generated as part of the PDB validation reports.
